# An Improved Search Algorithm to Find G-Quadruplexes in Genome Sequences

**DOI:** 10.1101/001990

**Authors:** Anna Varizhuk, Dmitry Ischenko, Igor Smirnov, Olga Tatarinova, Vyacheslav Severov, Roman Novikov, Vladimir Tsvetkov, Vladimir Naumov, Dmitry Kaluzhny, Galina Pozmogova

## Abstract

A growing body of data suggests that the secondary structures adopted by G-rich polynucleotides may be more diverse than previously thought and that the definition of G-quadruplex-forming sequences should be broadened. We studied solution structures of a series of naturally occurring and model single-stranded DNA fragments defying the G_3+_N_L1_G_3+_N_L2_G_3+_N_L3_G_3+_ formula, which is used in most of the current GQ-search algorithms. The results confirm the GQ-forming potential of such sequences and suggest the existence of new types of GQs. We developed an improved (broadened) GQ-search algorithm (http://niifhm.ru/nauchnye-issledovanija/otdel-molekuljarnoj-biologii-i-genetiki/laboratorija-iskusstvennogo-antitelogeneza/497-2/) that accounts for the recently reported new types of GQs.

## INTRODUCTION

Non-canonical polynucleotide structures play an important role in biogenesis processes, such as transcription, DNA repair, replication, translocation and RNA splicing (Saini et al. 2013). A clear view of DNA/RNA secondary structures and dynamics is necessary to understand the mechanisms of genomic regulation and to identify new biomarkers of pathology and drug targets. A growing body of data suggests that the secondary structures adopted by G-rich polynucleotides may be more diverse than previously thought (Kaluzhny et al. 2009; Tomasko et al. 2009; Guedin et al. 2010; Amrane et al. 2012; Beaudoin et al. 2013; Mukundan and Phan 2013). For instance, two new types of G-quadruplexes (GQs) have recently been reported: GQs with mismatches (Tomasko et al. 2009) and GQs with bulges (Mukundan and Phan 2013) (Figure 1). GQs with bulges (bGQs) are GQs in which two stacked tetrad-forming guanosines in one column are separated by a projecting nucleoside. GQs with mismatches (mGQs) contain one or more substitutions of G for other nucleotides in the tetrads. (The mismatching nucleosides may participate in stacking). Both types of structures appeared to be stable under physiological conditions. These findings have led the researchers to the conclusion that the definition of GQ-forming sequences should be broadened. All currently available online search tools for GQs (Quad finder (Scaria et al. 2006), QGRS Mapper (Kikin et al. 2006) and QGRS predictor (Menendez et al. 2012)) employ the G_3_+N_L1_G_3_+N_L2_G_3_+N_L3_G_3_+ formula, which only defines canonical (‘perfect’) GQs.

**Figure 1.**
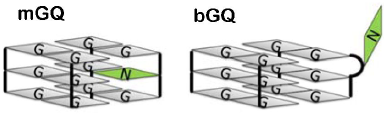
Imperfect GQ structures (imGQs). For each of the two imGQ types, a single example is shown. However, both mGQ and bGQ structures are diverse and can theoretically contain more than one mismatch or bulge. Loops are not shown.

We present here the first GQ-search tool, imGQfinder, that accounts for noncanonical (‘imperfect’) quadruplex structures (imGQs; i.e., bGQs and mGQs) in addition to canonical GQs. The ImGQfinder tool is freely accessible at the URL http://niifhm.ru/nauchnye-issledovanija/otdel-molekuljarnoj-biologii-i-genetiki/laboratorija-iskusstvennogo-antitelogeneza/497-2/. Structural studies of a series of (G_3+_N_L1_G_3+_N_L2_G_3+_N_L3_G_3+_)-defying oligonucleotides (ONs), whose imGQ-forming potential was predicted by imGQfinder, were performed to verify our improved GQ-search algorithm. We also utilize ImGQfinder for statistical analysis of the imGQ-motif distribution in the human genome.

## RESULTS

### ImGQ-motif definition and ImGQfinder interface (algorithm implementation)

In our broadened algorithm, we search for G-runs, determine the distance between them and select fragments that comply with the predetermined conditions for the maximum length of GQ loops and the minimum number of nucleotides in a G-run (i.e., the number of tetrads). The imGQ motif definition for imGQs with single defects is presented in Table 1. ImGQs with multiple defects can also be analyzed, but the relative set of formulas is not shown. Apparently, most imGQ motifs can be interpreted as both putative bGQs and mGQs. Some imGQs may also turn out to be ‘perfect’ GQs with fewer tetrads, e.g., a putative 3-tetrad imGQ with a bulge or a mismatch in the external tetrad can theoretically adopt the canonical 2-tetrad GQ conformation. ImGQfinder searches for all GQ and imGQ motifs, including overlapping ones. The program is implemented in Perl. The graphical user interface was developed using the Tk library. The inputs include the queried nucleotide sequence in fasta format, the number of tetrads and defects and the maximum loop length. The hits are displayed in a table. The coordinates of each G-run start (G1, G2, G3 and G4) and the positions of the defects (DEFECT) in each imGQ-forming fragment are shown. The user can also see the full sequences of the putative GQs or imGQs (the ‘Add sequence to output’ option).

**Table 1.** Monomolecular imGQ motifs with single defects (mismatches or bulges).

| Position of a mismatch/bulge | Formula |
| --- | --- |
| 5' –terminal G-run | $G_{i-1}NG_{k-i}(L_{xA}G_k)_3, A=1,2,3$ |
| The second G-run from the 5'-terminus | $G_k L_{x1} G_{i-1}NG_{k-i}(L_{xA}G_k)_2, A=2,3$ for mGQs<br>$G_k L_{x1} G_{i-1}NG_{k-i+1}(L_{xA}G_k)_2, A=2,3$ for bGQs |
| The third G-run from the 5'-terminus | $(G_k L_{xA})_2 G_{i-1}NG_{k-i} L_{x3}G_k, A=1,2$ for mGQs<br>$(G_k L_{xA})_2 G_{i-1}NG_{k-i+1} L_{x3}G_k, A=1,2$ for bGQs |
| 3'-terminal G-run | $(G_k L_{xA})_3 G_{i-1}NG_{k-i}, A=1,2,3$ |

L denotes any nucleotide in a loop; N denotes any mismatching (bulging) nucleotide (if N = G, the formula describes a canonical, 'perfect' GQ); A is the loop index; xA is the number of nucleotides in loop A; k is the number of guanosines in a G-run ( $k \geq 3$ ); and i is the position of a mismatch/bulge in the G-run ( $1 < i < k$ ).

In addition, we offer an application that can determine overlapping GQ/imGQ sites in the form of a single lengthy fragment with GQ/imGQ-forming potential (the ‘Add intersected output’ option). This feature is useful for estimating the maximum number of quadruplexes that can exist simultaneously.

### Structural studies (algorithm verification)

Although several recent publications contain direct evidence for the existence of imGQs that are stable under physiological conditions, such structures are still relatively new, and there are few examples of well-characterized imGQs. To complement the studies on imGQ structures and to verify our search algorithm for imGQs, we synthesized a set of naturally occurring and model single-stranded DNA fragments that were defined by ImGQfinder to be putative imGQs and GQs, and we analyzed their conformations in solution using physicochemical methods. The ONs are listed in Table 2.

**Table 2.**
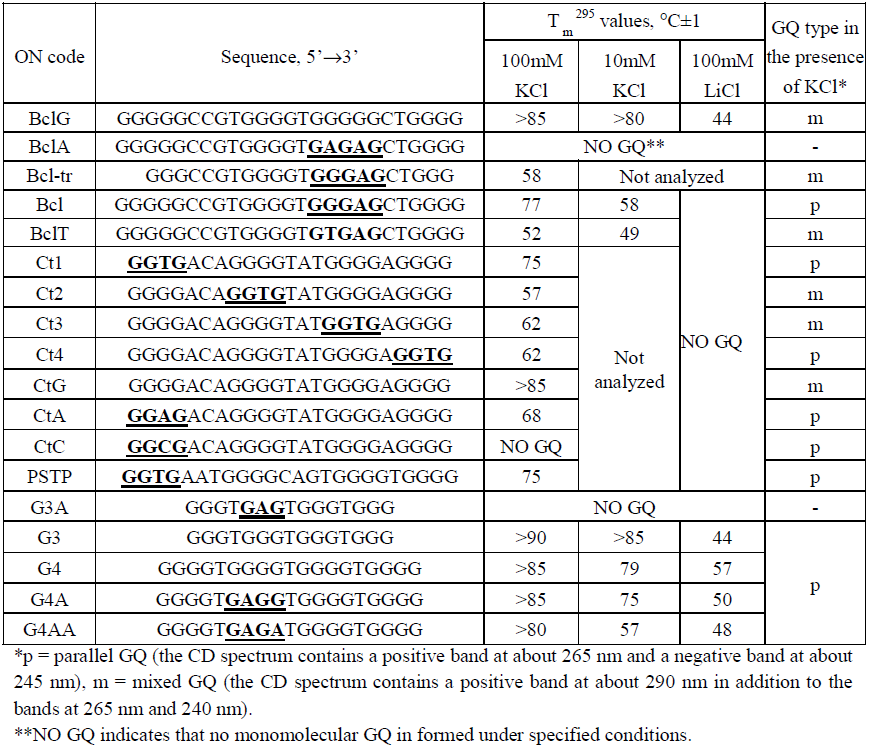
Sequences of GQ-forming ONs and characteristics of their solution structures. Pale pink indicates naturally occurring ON sequences. Pale blue indicates that no monomolecular GQ is formed under the specified conditions. Interrupted G3 and G4 runs are underlined.

The sequences Bcl, Ct1 and PSTP were taken from the human genome. Bcl is located in the BCL2 promoter region 42 nucleotides upstream of the translation start site (NCBI Reference Sequence: NC_000018.9, chr18: -60985942 to -60985966). Ct1 is located in the intron of the CTIF gene (NCBI Reference Sequence: NC_000018.9, chr18: +46379322 to +46379344). PSTP is located at the PSTPIP2 intron/boundary (NCBI Reference Sequence: NC_000018.9, chr18: +43572049 to +43572072). BclG, BclA, BclT, Bcl-tr (truncated), Ct2, Ct3, Ct4, CtA, CtC and CTG are mutants of Bcl and Ct1. G3, G3A, G4, G4A and G4AA are model sequences. The solution structures of the ONs were investigated using UV-melting experiments, CD spectroscopy and NMR spectroscopy. The rotational relaxation times of EtBr in complex with the ONs are proportional to the hydrodynamic volumes of the molecules, and these times were estimated to distinguish between monomolecular and intermolecular quadruplexes. The melting temperatures of monomolecular GQs/imGQs and the GQ characteristics determined from the CD data (parallel, antiparallel or mixed GQ folding) are given in Table 1. Fragments of the ^1^H-NMR spectra and CD spectra of the ONs Bcl, Ct1 and their mutants are shown in Figure 2. For the UV-melting profiles, molecularity analysis, CD spectra of the model ONs G3, G4 and their mutants and all the corresponding experimental procedures, see the supporting information.

**Figure 2.**
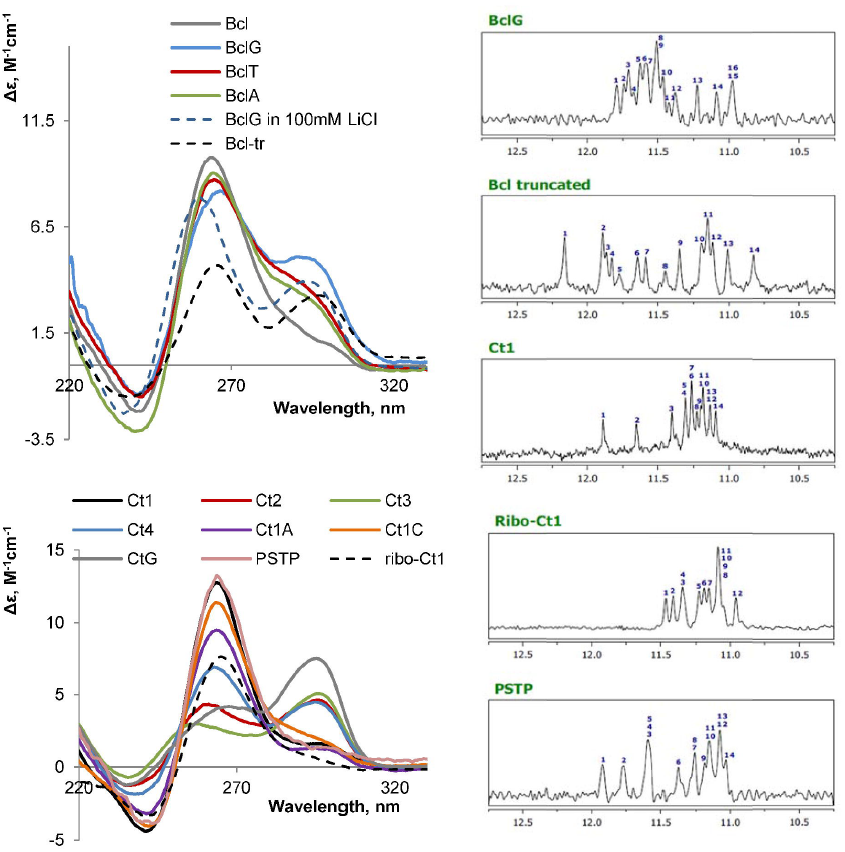
Left: CD spectra of the ONs Bcl, CT1 and their mutants. The ellipticity is given per mole of nucleotide. Buffer conditions: 100 mM KCl in 25 mM Tris-HCl (pH 7.5). Right: ^1^H NMR spectra fragments of several GQ- and imGQ-ONs. Buffer conditions: 100 mM KCl in 25 mM Tris-HCl (pH 7.5). ON concentration: 0, 1 mM.

The ONs G4, G3, CtG and BclG can fold into perfect 4-tetrad (G4, CtG and BclG) and 3-tetrad (G3) GQs according to the conventional GQ definition. Indeed, all of them formed highly stable parallel (G3^1^, G4 and CTG) or mixed (BclG) GQs in the presence potassium salt, as evidenced by the CD spectra and the UV-melting profiles. The imino region of the BclG ^1^H-NMR spectrum (Figure 2) contains 16 signals, which is consistent with 4 G-tetrads.

The ONs BclT, G4A, G4AA, Ct1-Ct4, CtA, CtC and PSTP defy the conventional G_3+_N_L1_G_3+_N_L2_G_3+_N_L3_G_3+_ formula and would be omitted by the currently existing GQ-search algorithms. ImGQfinder defines all of these sequences to be putative imGQs. Indeed, all these ONs form stable monomolecular quadruplexes in the presence of potassium salt. Fourteen signals in the imino regions of the ^1^H-NMR spectra of the ONs Ct1 and PSTP are consistent with 4-tetrad mGQ structures with one imperfect tetrad^2^. Twelve signals in the imino-spectrum region of ribo-Ct1 most likely suggest a 3-tetrad bGQ structure (Molecular modeling studies were performed to clarify the Ct1 structure. For more information, see the supporting data). Importantly, the ONs BclT and G4aa cannot even form 2-tetrad GQs according to the conventional GQ definition. However, these ONs appear to fold into rather stable GQ-like structures under physiological conditions. These results confirm that ImGQfinder can be used to predict the possibility of bGQ and mGQ formation.

### ImGQs in the human genome: statistical analysis (algorithm application)

ImGQfinder was utilized to reassess the abundance and to analyze the distribution of putative quadruplex sites in the human genome. We only considered 4-tetrad GQs and imGQs, which are generally more stable than 2- and 3-tetrad GQs according to the literature and our own physicochemical data. The sequences representing overlapping GQ/imGQ sites were counted only once (this was performed using an application feature of ImGQfinder). Sites with both GQ- and imGQ-folding potentials were regarded as putative GQs because the latter are generally more stable. As expected, imGQs are substantially more abundant than GQs (Table 3). Thus, the maximum overall number of quadruplex-like structures realized in vivo may be significantly higher than previously thought. The distribution of putative imGQs and GQs within RefSeq genes (genomic sequences used as reference standards for well-characterized genes; http://www.ncbi.nlm.nih.gov/refseq/rsg/about/) is shown in Figure 3 (for a more detailed analysis, see the supporting information). As expected, imGQs are substantially more abundant than GQs.

**Table 3.**
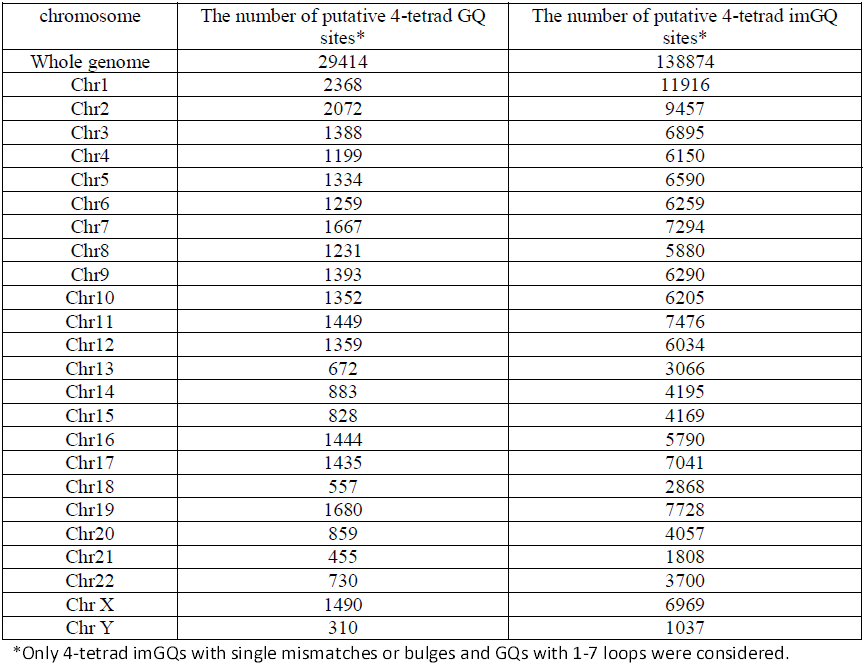
GQ and imGQ abundance in human genome (distribution among chromosomes).

| chromosome | The number of putative 4-tetrad GQ sites* | The number of putative 4-tetrad imGQ sites* |
| --- | --- | --- |
| Whole genome | 29414 | 138874 |
| Chr1 | 2368 | 11916 |
| Chr2 | 2072 | 9457 |
| Chr3 | 1388 | 6895 |
| Chr4 | 1199 | 6150 |
| Chr5 | 1334 | 6590 |
| Chr6 | 1259 | 6259 |
| Chr7 | 1667 | 7294 |
| Chr8 | 1231 | 5880 |
| Chr9 | 1393 | 6290 |
| Chr10 | 1352 | 6205 |
| Chr11 | 1449 | 7476 |
| Chr12 | 1359 | 6034 |
| Chr13 | 672 | 3066 |
| Chr14 | 883 | 4195 |
| Chr15 | 828 | 4169 |
| Chr16 | 1444 | 5790 |
| Chr17 | 1435 | 7041 |
| Chr18 | 557 | 2868 |
| Chr19 | 1680 | 7728 |
| Chr20 | 859 | 4057 |
| Chr21 | 455 | 1808 |
| Chr22 | 730 | 3700 |
| Chr X | 1490 | 6969 |
| Chr Y | 310 | 1037 |

\*Only 4-tetrad imGQs with single mismatches or bulges and GQs with 1-7 loops were considered.

**Figure 3.**
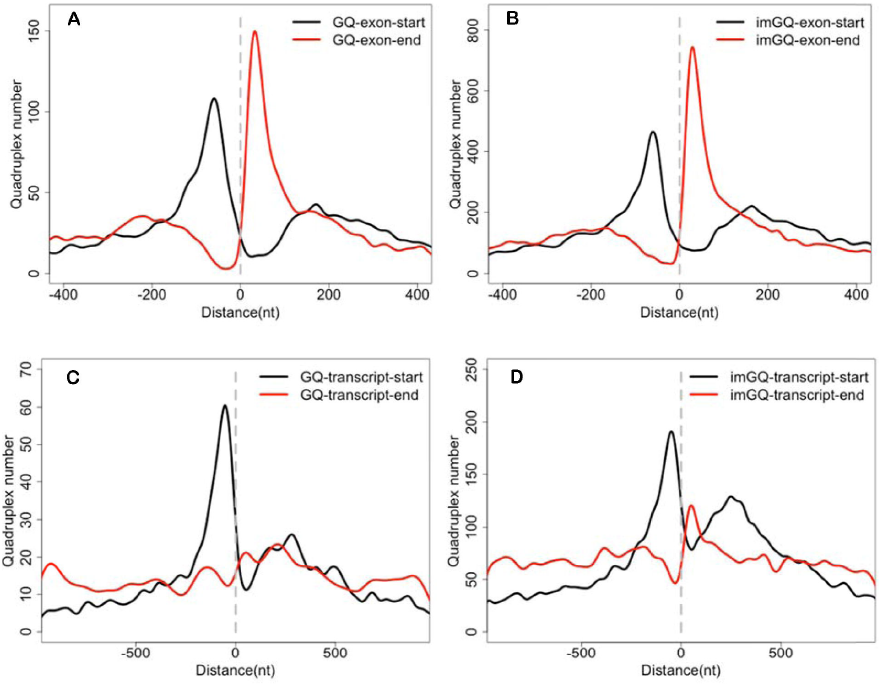
GQ and imGQ distribution within genes. A: GQ locations relative to exon/intron boundaries; B: imGQ locations relative to exon/intron boundaries; C: GQ locations relative to transcription start sites; D: imGQ locations relative to transcription start sites.

To additionally validate the ImGQfinder algorithm, we also calculated the number of all putative non-overlapping ‘perfect’ 3-tetrad guadruplexes in the human genome and compared it with the literature data. The obtained value (359 k) is close to the previous estimations (376 k) (Huppert and Balasubramanian 2005).

## DISCUSSION

A new GQ-search algorithm, which is based on a broadened definition of quadruplex-forming sequences, and the user-friendly online tool ImGQfinder were developed. The algorithm was verified by structural studies of a series of ONs whose imGQ-forming potential was predicted by imGQfinder. Importantly, the physicochemical properties of ImGQs and GQ, such as the thermal stability under physiological conditions, appear to be rather similar.

Reassessment of the abundance of putative quadruplex sites in the human genome with imGQfinder revealed that the maximum number of G4 structures that could be simultaneously realized has been underestimated. As is evident from Figure 3, putative GQ and imGQ sites have basically similar distributions within RefSeq genes. Exons tend to be depleted of both GQs and imGQs. Large clusters of putative GQ/imGQ sites were found in the introns near the intron/exon boundaries and in the promoters that are approximately 100 bp downstream of the transcription start site. GQ clustering in 5’ untranslated regions is consistent with literature data (Huppert and Balasubramanian 2007; Maizels and Gray 2013). The results of several recent studies suggest 5’-UTR GQ participation in transcription and translational regulation (Huppert et al. 2008). GQs at intron/exon boundaries may play a role in splicing. Although known enhancer/silencer splicing element motifs (Wang et al. 2005) do not have GQ/imGQ-folding potential, recent publications suggest that GQ-like structures may influence splicing (Han et al. 2005; Fisette et al. 2012) and that the genes that undergo alternative splicing are enriched with GQs (Kostadinov et al. 2006).

In conclusion, the broadened GQ-search algorithm opens up new opportunities in the prediction of DNA/RNA structure and allows thorough analysis of all possible conformations adopted by polynucleotides.

## METHODS

**The ON synthesis and purification, the MS analysis and the UV-melting, CD and rotational relaxation time measurements** were performed as previously described (Varizhuk et al. 2013).

For the **analysis GQ/imGQ abundance and distribution** in human genome, RefSeq genomic sequences (http://www.ncbi.nlm.nih.gov/refseq/rsg/about/) were used.

### NMR studies

NMR samples were prepared at a concentration of ∼0.1 mM in 0.6 ml H2O+D2O (10%) buffer solution containing 20 mM Tris-HCl (pH 7.5) and 100 mM KCl and annealed (heated to 90 C for 3 minutes, then cooled quickly on ice) prior to spectral measurements to ensure unimolecular quadruplex folding. ^1^H-NMR spectra were recorded with Bruker AVANCE II 300 (300.1 MHz), Bruker AMX III (400.1 MHz) and Bruker AVANCE II 600 (600.1 MHz) spectrometers. The ^1^H chemical shifts were referenced relative to an external standard - sodium 2, 2-dimethyl-2-silapentane-5-sulfonate (DSS). The spectra were recorded using presaturation or pulsed-field gradient WATERGATE W5 pulse sequences (zgprsp and zggpw5 from the Bruker library, respectively) for H_2_O suppression.

### Molecular modeling

Ct1 GQ models 1 and 2 were created as follows. The starting positions of the GQ core atoms were obtained from the PDB (139D and 2KQH). The core of every GQ was created using Swiss-PDB Viewer. Then, loops were added step by step as described further by utilizing the SYBYL 8.0 molecular modeling package. To remove unfavorable van der Waals interactions, the models were reoptimized after attaching each loop using SYBYL 8.0 and the Powell method with the following parameters: Gasteiger-Hückel charges, TRIPOS force field, non-bonded cut-off distance of 8 Ǻ, distance-dependent dielectric function, 1000 iterations, the simplex method in an initial optimization and the energy gradient convergence criterion with a threshold of 0.05 kcal*mol^-1^*Å^-1^. The GQ core was frozen during the above reoptimizations.

The molecular dynamics simulations (MD) were performed with the Amber 9 suite with ff99SB and parmbsc0 force fields as described previously (Varizhuk et al. 2013). The trajectory length was 35 ns. Snapshot visualization and hydrogen bond analysis were performed using VMD (http://www.ks.uiuc.edu/Research/vmd/) with a donor-acceptor distance of 3 Å and an angle cutoff of 20 degrees. Snapshots were taken every 0.1 ns.

The MM-GBSA method was used to calculate the Ct1 GQ free energies for models 1 and 2. In this approach, the free energy is calculated according to the formula *G* = *E*_*MM*_ + *G*_*sol*_ −*TS*, where *E*_*MM*_, *G*_*sol*_ and *TS* are the total mechanical energy of the molecule in gas phase, the free energy of hydration and the entropic contribution, respectively. *E*_*MM*_ was calculated as the sum of the electrostatic energies, van der Waals energies and the energies of internal strain (bonds, angles and dihedrals) by using a molecular-mechanics approach. *G*_*sol*_ was calculated as the sum of the polar (*G*_*polar*_) and nonpolar (*G*_*nonpolar*_) terms. The electrostatic contribution to the hydration energy *G*_*polar*_ was computed using the Generalized Born (GB) method (Onufriev et al. 2000) using the algorithm developed by Onufriev et al. (Weiser et al. 1999; Onufriev et al. 2002) for calculating the effective Born radii. The non-polar component of the hydration energy *G*_*nonpolar*_, which includes solute-solvent van der Waals interactions and the free energy of cavity formation in solvent, was calculated using the following formula: *G*_*nonpolar*_ = *α* * *SASA*, where SASA is the solvent accessible surface area. SASA was computed using the LCPO method (Srinivasan et al. 1998) with α = 0.00542 kcal/mol^-1^ Å^-2^. The entropic term was not calculated explicitly, but it was accounted for implicitly via the GQ conformational mobility. Snapshots taken from a single trajectory of the MD simulation of the complex were used for the calculations of the binding free energy.

## DATA ACCESS

ImGQfinder is freely accessible at the URL http://niifhm.ru/nauchnye-issledovanija/otdel-molekuljarnoj-biologii-i-genetiki/laboratorija-iskusstvennogo-antitelogeneza/497-2/.

## ACKNOWLEDGEMENTS

The work has been supported by Russian Foundation for Basic Research [14-04-01244] and the program of the Presidium of the Russian Academy of Sciences on Molecular and Cell Biology. We thank A. Aseychev for his help with thrombin time measurements and V. Karpov for oligonucleotide synthesis.

### DISCLOSURE DECLARATION

We declare no conflict of interest.

## Footnotes

1 G3 demonstrated extreme stability in potassium, and the stability was even superior to that of G4. We attribute this stability to the fact that single-nucleotide fragments separating G runs fit perfectly well in the diagonal loops of 3-tetrad GQs but may be slightly too short for 4-tetrad diagonal loops.

2 One imperfect tetrad contains three Hoogsteen-bound Gs with two imino G protons that participate in H-bonding, which results in two additional signals in the relative region of the ^1^H-NMR spectrum.

## REFERENCES

Amrane S, Adrian M, Heddi B, Serero A, Nicolas A, Mergny JL, Phan AT. 2012. Formation of pearl-necklace monomorphic G-quadruplexes in the human CEB25 minisatellite. J Am Chem Soc 134(13): 5807–5816.

Beaudoin JD, Jodoin R, Perreault JP. 2013. New scoring system to identify RNA G-quadruplex folding. Nucleic Acids Res.

Fisette JF, Montagna DR, Mihailescu MR, Wolfe MS. 2012. A G-rich element forms a G-quadruplex and regulates BACE1 mRNA alternative splicing. J Neurochem 121(5): 763–773.

Guedin A, Gros J, Alberti P, Mergny JL. 2010. How long is too long? Effects of loop size on G-quadruplex stability. Nucleic Acids Res 38(21): 7858–7868.

Han K, Yeo G, An P, Burge CB, Grabowski PJ. 2005. A combinatorial code for splicing silencing: UAGG and GGGG motifs. PLoS Biol 3(5): e158.

Huppert JL, Balasubramanian S. 2005. Prevalence of quadruplexes in the human genome. Nucleic Acids Res 33(9): 2908–2916.

Huppert JL, Balasubramanian S. 2007. G-quadruplexes in promoters throughout the human genome. Nucleic Acids Res 35(2): 406–413.

Huppert JL, Bugaut A, Kumari S, Balasubramanian S. 2008. G-quadruplexes: the beginning and end of UTRs. Nucleic Acids Res 36(19): 6260–6268.

Kaluzhny D, Shchyolkina A, Livshits M, Lysov Y, Borisova O. 2009. A novel intramolecular G-quartet-containing fold of single-stranded d(GT)(8) and d(GT)(16) oligonucleotides. Biophys Chem 143(3): 161–165.

Kikin O, D'Antonio L, Bagga PS. 2006. QGRS Mapper: a web-based server for predicting G-quadruplexes in nucleotide sequences. Nucleic Acids Res 34(Web Server issue): W676–682.

Kostadinov R, Malhotra N, Viotti M, Shine R, D'Antonio L, Bagga P. 2006. GRSDB: a database of quadruplex forming G-rich sequences in alternatively processed mammalian pre-mRNA sequences. Nucleic Acids Res 34(Database issue): D119–124.

Maizels N, Gray LT. 2013. The G4 genome. PLoS Genet 9(4): e1003468.

Menendez C, Frees S, Bagga PS. 2012. QGRS-H Predictor: a web server for predicting homologous quadruplex forming G-rich sequence motifs in nucleotide sequences. Nucleic Acids Res 40(Web Server issue): W96–W103.

Mukundan VT, Phan AT. 2013. Bulges in G-Quadruplexes: Broadening the Definition of G-Quadruplex-Forming Sequences. J Am Chem Soc.

Onufriev A, Bashford D, Case DA. 2000. Modification of the Generalized Born Model Suitable for Macromolecules. J Phys Chem B 104(15): 3712–3720.

Onufriev A, Case DA, Bashford D. 2002. Effective Born radii in the generalized Born approximation: the importance of being perfect. J Comput Chem 23(14): 1297–1304.

Saini N, Zhang Y, Usdin K, Lobachev KS. 2013. When secondary comes first–the importance of non-canonical DNA structures. Biochimie 95(2): 117–123.

Scaria V, Hariharan M, Arora A, Maiti S. 2006. Quadfinder: server for identification and analysis of quadruplex-forming motifs in nucleotide sequences. Nucleic Acids Res 34(Web Server issue): W683–685.

Srinivasan J, Miller J, Kollman PA, Case DA. 1998. Continuum solvent studies of the stability of RNA hairpin loops and helices. J Biomol Struct Dyn 16(3): 671–682.

Tomasko M, Vorlickova M, Sagi J. 2009. Substitution of adenine for guanine in the quadruplex-forming human telomere DNA sequence G(3)(T(2)AG(3))(3). Biochimie 91(2): 171–179.

Varizhuk AM, Tsvetkov VB, Tatarinova ON, Kaluzhny DN, Florentiev VL, Timofeev EN, Shchyolkina AK, Borisova OF, Smirnov IP, Grokhovsky SL et al. 2013. Synthesis, characterization and in vitro activity of thrombin-binding DNA aptamers with triazole internucleotide linkages. Eur J Med Chem 67C: 90–97.

Wang J, Smith PJ, Krainer AR, Zhang MQ. 2005. Distribution of SR protein exonic splicing enhancer motifs in human protein-coding genes. Nucleic Acids Res 33(16): 5053–5062.

Weiser J, Shenkin PS, Still WC. 1999. Approximate Atomic Surfaces from Linear Combinations of Pairwise Overlaps (LCPO). J Comput Chem 20: 217–230.

